# A Hall-Effect Sensor-Based Queen Bee Detection System – a Proof of Concept

**DOI:** 10.64898/2026.06.29.735221

**Authors:** Bartosz Krajnik, Monika Maciejewska, Stanisław Janeczko, Andrzej Szczurek

## Abstract

The queen bee is the central individual responsible for colony establishment, growth, and survival. Reliable confirmation of successful mating, continued queen presence, and normal reproductive performance is essential for effective colony management. We present a queen bee detection system based on an array of Hall-effect sensors and a miniature magnetic tag attached to the queen. The system is designed for continuous operation and real-time monitoring. A prototype was developed, constructed, and evaluated under both laboratory and field conditions. Field experiments conducted in an apiary demonstrated that the system can reliably detect queen bee passages through the hive entrance, enabling the identification of activities associated with mating flights. The results confirm the feasibility of Hall-effect sensing for automated, non-invasive queen bee monitoring and establish magnetic sensing as a promising new measurement modality for precision apiculture.

## 1. Introduction

Honey bees are indispensable to both natural ecosystems and human well-being. As major pollinators, they play a fundamental role in maintaining biodiversity and sustaining agricultural production, thereby contributing directly to global food security. Preserving healthy and viable honey bee populations is therefore of considerable societal importance. Consequently, research into honey bee biology, colony dynamics, and sustainable beekeeping practices remains essential for ensuring ecosystem resilience and long-term food production.

Within a honey bee colony, the queen is the sole reproductive female and the central individual responsible for colony establishment, growth and survival. A colony deprived of its queen inevitably declines unless it successfully rears a replacement queen or a new queen is introduced by the beekeeper. Throughout most of her life, the queen remains inside the hive, where her primary biological function is egg laying. This reproductive activity becomes possible only after successful mating, a process that determines the genetic composition of the entire colony because both maternal and paternal genomes are transmitted to subsequent generations. Consequently, mating success influences numerous colony traits associated with productivity, behaviour and resilience, including resistance to pathogens and parasites such as *Varroa destructor* (Martin et al., 2024). Reliable confirmation of successful mating, continued queen presence and normal reproductive performance is therefore fundamental to colony development and long-term survival.

In current beekeeping practice, queen presence is typically verified through manual colony inspections. These inspections are labour-intensive, disruptive to colony activity and require substantial practical experience, while even skilled beekeepers may occasionally fail to locate the queen. Moreover, routine inspections are discouraged during the mating period, when disturbance may interfere with normal queen behaviour. Under typical conditions, the mating period extends for approximately three weeks following queen emergence. During this time, the queen performs one or more mating flights to drone congregation areas, where polyandrous mating with multiple drones promotes genetic diversity within honey bee populations (Büchler et al., 2025). Preserving these natural mating processes is important not only for maintaining colony fitness but also for commercial queen breeding, as naturally mated queens generally exhibit higher breeding value and consequently greater economic value.

The importance of continuous information on queen status, together with the practical limitations of manual inspections, has stimulated the development of automated monitoring technologies capable of providing non-invasive and continuous observations (Anuar et al., 2023). Such systems constitute an important component of emerging precision apiculture, enabling data-driven colony management and supporting intelligent decision-support systems (Hadjur et al., 2022).

Current approaches to automated queen monitoring are based on two main sensing principles. The first relies on analysis of acoustic signals generated by the colony. Previous studies have identified queen-specific acoustic signatures and characteristic behavioural sounds associated with queen presence, queenlessness and swarming (Uthoff et al., 2023). Spectral analysis combined with artificial neural networks has been employed to classify colony status (Kanelis et al., 2023; Ruvinga et al., 2023), while the characteristic piping signals emitted by virgin queens have been used to detect swarming-related events (Rybin et al., 2017). Although acoustic monitoring is entirely non-invasive and provides valuable information on colony dynamics, reliable interpretation requires sophisticated signal-processing algorithms and extensive reference datasets.

The second approach employs radio-frequency identification (RFID). RFID technology has been successfully applied to record queen departures from and returns to the hive, particularly in studies investigating mating-flight behaviour (Heidinger et al., 2014). Recent developments have enabled the use of miniature RFID tags only a few millimetres in size, allowing reliable identification at distances of approximately 1–2 cm when the reader is integrated into the hive entrance (de Souza et al., 2018). While RFID provides highly reliable event detection, its effective operating range is inherently limited by the characteristics of the technology (Alburaki et al., 2021).

Despite the considerable progress achieved with acoustic and RFID-based monitoring, alternative sensing modalities remain largely unexplored. In particular, magnetic sensing has not previously been investigated as a method for automated queen detection.

In the present study, we introduce a queen detection system based on an array of Hall-effect sensors and a miniature magnetic tag attached to the queen. A prototype device was designed, constructed and evaluated under both laboratory and field conditions. The obtained results demonstrate the feasibility of Hall-effect sensing for automated, non-invasive queen detection and establish magnetic sensing as a promising new measurement modality for precision apiculture. To the best of our knowledge, this study is the first to report the application of Hall-effect sensor technology for honey bee queen monitoring.

## 2. Detection system

### 2.1 Detection principle

Our detection system uses the Hall Effect as the detection principle. When current flows through a conductive plate placed in a perpendicular magnetic field, the Lorentz force deflects the charge carriers to one side. This creates a measurable “Hall voltage” across the plate, which is directly proportional to the magnetic field’s strength. A Hall sensor operates according to the Hall Effect.

We proposed detecting the queen bee tagged with a miniature magnet by an array of Hall sensors.

### 2.2 System architecture and components

The detection system consists of three modules: a sensing module, a data acquisition, processing, storage, and communication module, and a power supply module. The overall architecture of the detection system is shown in Fig. 1.

**Fig. 1.**
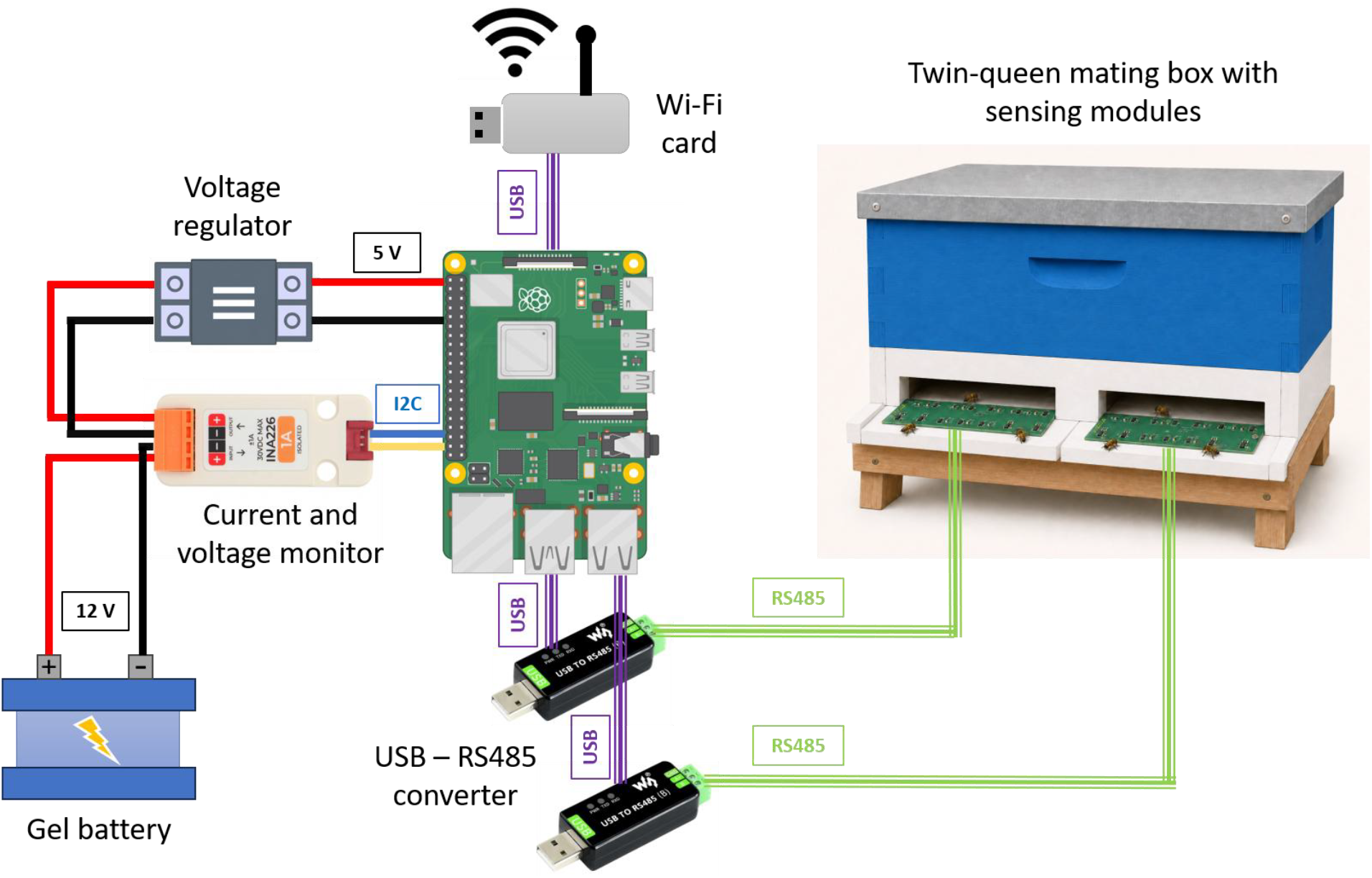
Diagram of the detection system.

**Fig. 2.**
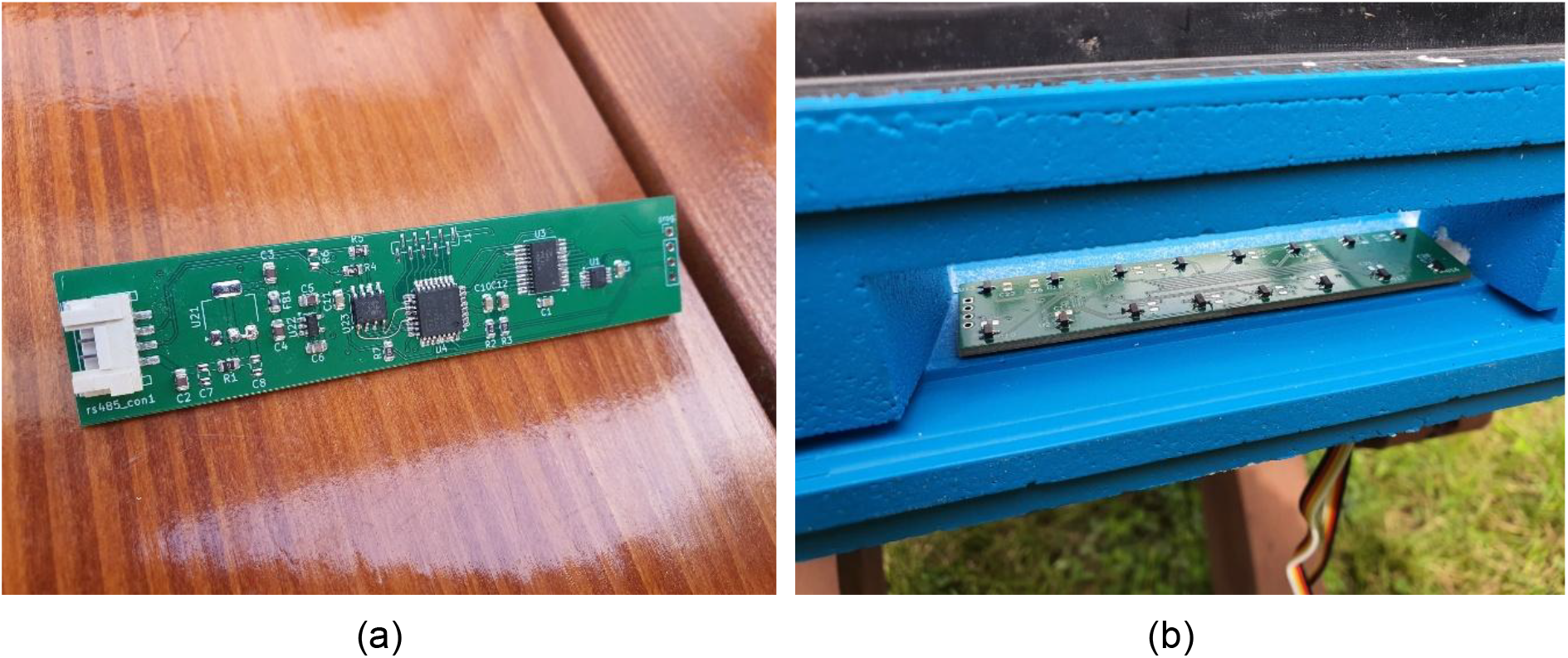
Dedicated PCB designed for sensor module utilising Hall-effect sensors. (a) Top of the PCB as manufactured. (b) Bottom side of the PCB mounted in the beehive entrance. Two rows of Hall-effect sensors are shown, eight sensors each.

#### Sensing module

The detection system employs miniature Hall-effect sensors. The Hall-effect sensor detects the magnetic field generated by a permanent magnet in its vicinity. The DRV5055 Ratiometric Linear Hall-Effect Sensor (Hall-effect sensor, 2026) was selected. The sensor responds proportionally to magnetic flux density. The system employs both 3.3 V and 5 V supply rails. When no magnetic field is present, the analog output drives half of the supply voltage (VCC). The output changes linearly with the applied magnetic flux density, and various sensitivity options enable maximal output voltage swing based on the required sensing range. North and south magnetic poles produce positive and negative shifts in the output voltage.

In the proposed solution, the magnetic field sensors are arranged in the form of an array. The sensors are mounted on the bottom side of a custom-designed two-layer PCB and organized into two rows, each consisting of eight Hall-effect sensors. The top side of the PCB accommodates the remaining electronic components, including an analog multiplexer, an analog-to-digital converter (ADC), a microcontroller, and a voltage regulator. The dedicated PCB enables synchronized acquisition of the sensor outputs while maintaining a compact mechanical design suitable for installation at the hive entrance.

The sensing module was designed to be integrated with the beehive. It has been mounted on the bottom of the beehive entrance, with the sensors facing upward. The module is connected to the other modules of the sensor system via cables routed through the base of the hive.

#### Data acquisition, processing, storage, and communication module

The data acquisition, processing, storage, and communication module consists of a Raspberry Pi 4B microcomputer, RS-485-to-USB converters, and an external Wi-Fi adapter. Measurement data from the sensing module are continuously transmitted to the Raspberry Pi using the industrial RS-485 communication standard at a sampling rate of 30 samples per second. The incoming data are pre-processed by a dedicated software service, which averages consecutive groups of five samples, resulting in an effective sampling rate of 6 samples per second. The processed data are buffered in memory and written to an external USB storage device once every minute. In addition, the measurement data are automatically synchronized with a remote server every 10 minutes to facilitate data access and provide secure backup. Network connectivity is provided by an external Wi-Fi adapter equipped with an external antenna connector, ensuring a stable Internet connection under field conditions. Secure remote access to the measurement station is enabled through the WireGuard VPN service, allowing authorized users to access the system from any computer connected to the Internet.

#### Power supply module

According to the initial design assumptions, the total power consumption of the sensing system is approximately 4 W. The system is powered by a 12 V, 70 Ah sealed gel lead-acid battery, which provides autonomous operation for approximately 5–6 days and requires only periodic recharging. The battery capacity was selected to ensure that its size and weight allow convenient manual transportation without the use of a vehicle. The system design also provides easy access to the battery, allowing it to be recharged or replaced without interrupting the operation of the sensing system.

The battery voltage and current are continuously monitored using an INA226 current and power monitor, which communicates with the Raspberry Pi 4B via the I^2^C interface. This enables continuous supervision of the battery status as well as real-time monitoring of the system power consumption. The collected measurements provide the basis for estimating the remaining operating time before battery recharging is required.

#### Events and System status Communication

The system provides the user with two categories of information: (1) event detection: The event is defined as the sensor signal change exceeding the predefined threshold – potentially indicating queen departure from the hive or her return, and (2) system status information, including the status of the power supply (battery voltage and current draw) and the quality of the wireless network connection, including Wi-Fi signal strength.

### 2.3 Magnetic tag

We proposed to tag the queen bee with a magnetic tag to enable its detection by Hall-effect sensors.

Tagging insects for behavioral monitoring is not a novel concept and has also been widely applied in honey bee research. Various tagging methods, including paint markers, plastic discs, and electronic tags, have been employed to identify and track individual insects. In beekeeping practice, queen bees are routinely marked to facilitate age tracking and visual identification during hive inspections. For this purpose, color-coded or numbered opalith discs are typically attached to the dorsal surface of the thorax. This approach, however, supports only manual inspection. RFID technology, which has been extensively used for studying insect behavior, including that of honey bees, also relies on tagging. RFID tags incorporate miniature antennas that enable insect detection by RFID readers.

Tagging always involves a compromise between maximizing the utility of the tag for the detection system and minimizing interference with the normal functioning of the insect. The solution presented in this study similarly requires the queen bee to be tagged. We proposed using a permanent magnet, to enable queen detection by Hall-effect sensors. To achieve good detectability, a neodymium magnet was selected. It provides a strong magnetic field while maintaining while the size of magnet is relatively small. The selection of the remaining design parameters of the magnetic tag was guided by recommendations derived from previous studies on insect tagging. So far, concerns have been raised regarding the weight and dimensions of attached tags.

#### Weight Considerations

The body mass of a virgin queen bee typically ranges from 170 mg to 220 mg. Pilot experiments reported by Hayworth et al. (2009) demonstrated that an additional dorsal load of 70 mg represented the maximum weight with which queen bees were still capable of flight. Additional loads of 60 mg and 30 mg significantly reduced the number of flights, mean flight duration, and the overall time spent flying. Furthermore, mating success, measured by sperm quantity and the number of patrilines detected among offspring, was negatively correlated with the added weight. In contrast, the study presented by Costa et al. (2021) evaluated the influence of RFID tag weight on the flight performance of *Melipona seminigra* and found that flight capacity was affected primarily by landscape conditions rather than by the additional payload.

For comparison, conventional opalith tags typically weigh between 0.5 and 1.5 mg. The mass of commercially available RFID tags used in standard bee-monitoring applications generally remains below 5 mg (Dolasevic et al., 2025). However, Beltrán et al. (2024) and Lorenzo-Lopez and Juan-Llacer (2025) presented a prototype RFID antenna weighing approximately 25 mg designed for queen bee localization within the hive.

Based on these findings, a design criterion was adopted for the proposed detection system stating that the magnet mass should not exceed approximately 10% of the body mass of a virgin queen bee, i.e. less than 17–22 mg.

#### Size Considerations

The average dimensions of a queen bee are approximately 20–25 mm in length and 4 mm in thorax width. A standard opalith tag is a plastic disc approximately 2.5 mm in diameter and 0.5–1 mm thick. According to Alburaki et al. (2021), the optimal dimensions of RFID tags used for tracking queen mating flights are approximately 2.5 × 2.5 × 0.4 mm. Also smaller tags measuring 1 × 1.6 × 0.5 mm have been successfully employed in this application, when the RFID reader was installed at the hive entrance (Heidinger et al., 2014). Larger dimensions were required in the case of queen location from outside the hive, where the antenna measured 3.09 × 2.61 mm (Beltrán et al., 2024). Concerns were raised that tags of this size could interfere with the maneuverability of virgin queens during mating flights and with their movement on the comb (Heidinger et al., 2014). Based on these considerations, in our prototype it was assumed that no dimension of the magnetic tag should exceed approximately 50% of the thorax width of a virgin queen bee, corresponding to a maximum dimension of about 2 mm.

#### Tag Placement

In insect-monitoring applications, tags are most commonly attached to the thorax (Dolasevic et al., 2025). Studies using oil-based paint markers have demonstrated superior color retention on the thorax compared with the abdomen or wings. Thoracic placement is also advantageous because it does not interfere with oviposition. During egg laying, the queen inserts only her abdomen into the brood cell, while the head and thorax remain outside. Consequently, the magnetic tag in the proposed system was mounted on the dorsal surface of the thorax.

#### Attachment Method

Tag attachment in beekeeping applications is typically achieved using non-toxic adhesives or resin-based compounds. Commercial cyanoacrylate adhesives have demonstrated satisfactory performance in field studies (Beltrán et al., 2024). For the prototype system, and until a more specialized attachment method is developed, the magnetic tag was fixed using a procedure identical to that commonly employed by queen breeders for attaching opalith tags.

#### Durability and Operational Risks

Unlike worker bees, whose relatively short lifespan often necessitates continuous replacement of tagged individuals in experimental studies, queen bees typically remain in the colony for extended periods. Consequently, tag longevity is less problematic in queen-monitoring applications. Nevertheless, there remains a possibility of tag detachment due either to natural causes or to mechanical abrasion resulting from contact with hive structures and colony members. The same risk applies to magnetic tags. In the case of magnetic tagging, an additional risk arises from potential interactions with metallic items. Such interactions may result in unintended magnet detachment, temporary immobilization of the queen within the hive, or, in the worst case, loss of the queen if such an event occurs during a flight.

## 3. Field tests

### 3.1 Experimental stand

Field tests were performed to validate the performance of the prototype system for queen bee detection in the beehive entrance. Twin-queen mating boxes were chosen for the field experiment. They allow monitoring two queens in parallel, using one detection system fitted with two sensing modules. A complete experimental stand is shown in Fig. 5.

After confirming that the installed detection system was fully operational, the nucs were populated with small bee colonies while providing sugar syrup as feed. In parallel, the newly emerged queen bees were tagged with magnets. Tags were glued centrally onto the scutum of the queens’ thoraxes to avoid interference with wing movements, as shown in Fig. 3. The magnetic tag was a neodymium magnet N38, 1.5×1.5 mm (cylinder). The tagged queens were introduced into the nucs which remained closed. After three days, the hives’ entrances were opened to let queen bees leave for mating flights. The readings from the detection system were monitored continuously, expecting that successful mating should occur within three weeks after introducing the queen.

**Fig. 3.**
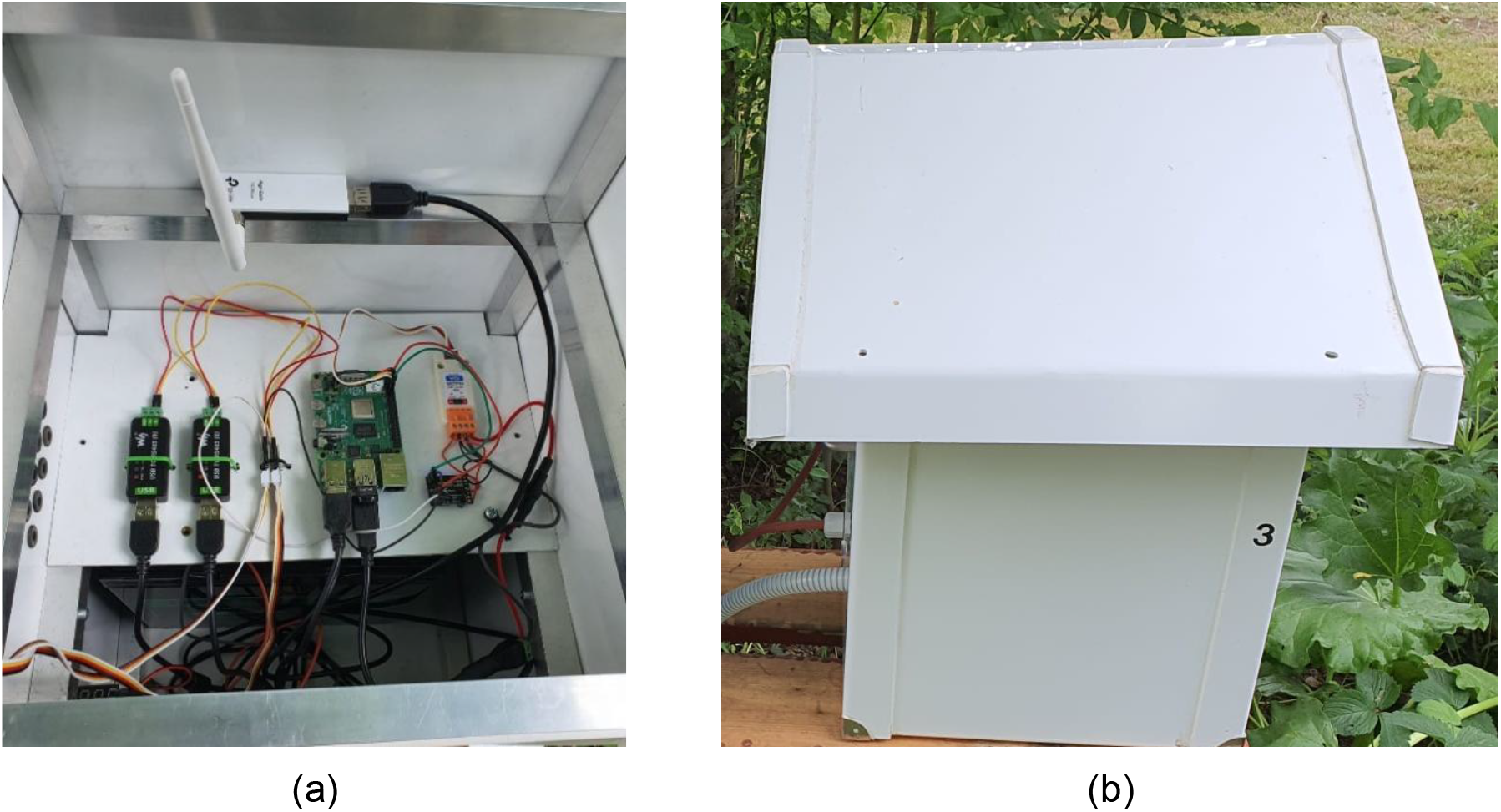
(a) Data acquisition, processing, storage, and communication module mounted on the upper shelf and the gel lead-acid battery mounted below the shelf, both enclosed within a self-contained casing. (b) Self-contained casing with both modules installed, providing protection against adverse weather conditions. External connectors are provided for the sensing module and battery charging.

**Fig. 4.**
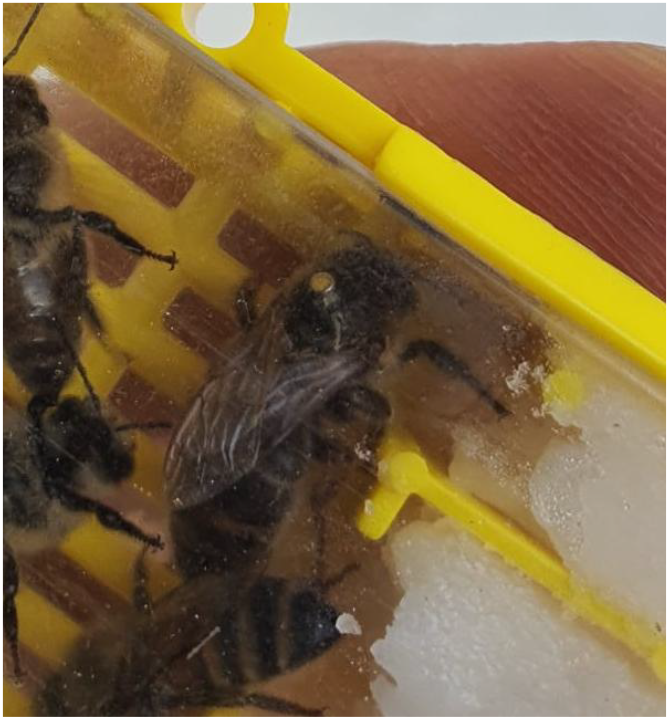
Queen bee tagged with a neodymium magnet N38, 1.5×1.5 mm.

**Fig. 5.**
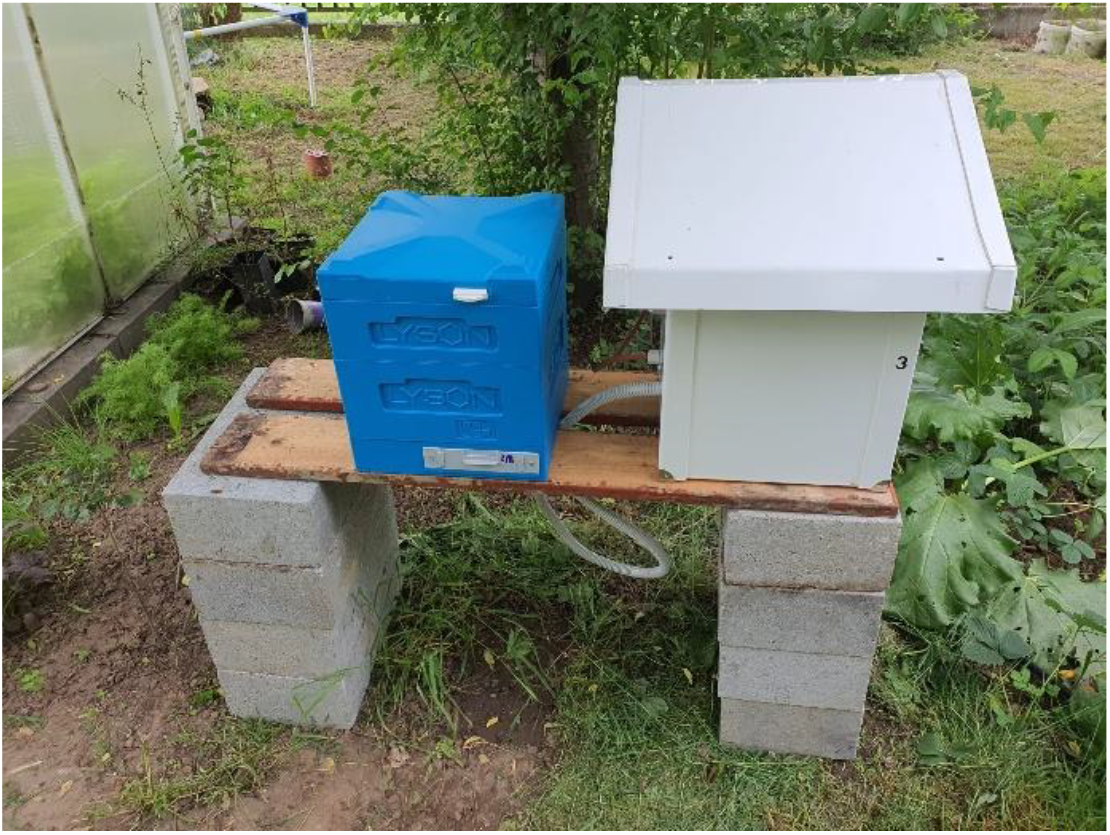
An experimental stand for testing, in field conditions, the queen bee detection system. Left: twin-queen mating box fitted with two sensing modules, one module per queen. Right: Housing with data acquisition, processing, storage, and communication module and a gel lead-acid battery inside.

### 3.2 Measurement results

In the following, we present the data collected during the experiment for which successful queen mating was confirmed. Beehive inspection revealed the queen bee tagged with the magnet and fresh eggs inside the comb cells.

Based on the measurement data, three categories of sensor signals were identified: no event, signals not associated with the queen bee detection, and signals potentially associated with queen bee detection. It should be noted that we refer to sensor signal, recorded over a period of time, at least for several minutes and not to a single sensor reading. Fig. 6 illustrates three identified categories of signals, which may be observed in a daily record of the measurement data from the detection system. The record includes the signals collected from eight channels.

**Fig. 6.**
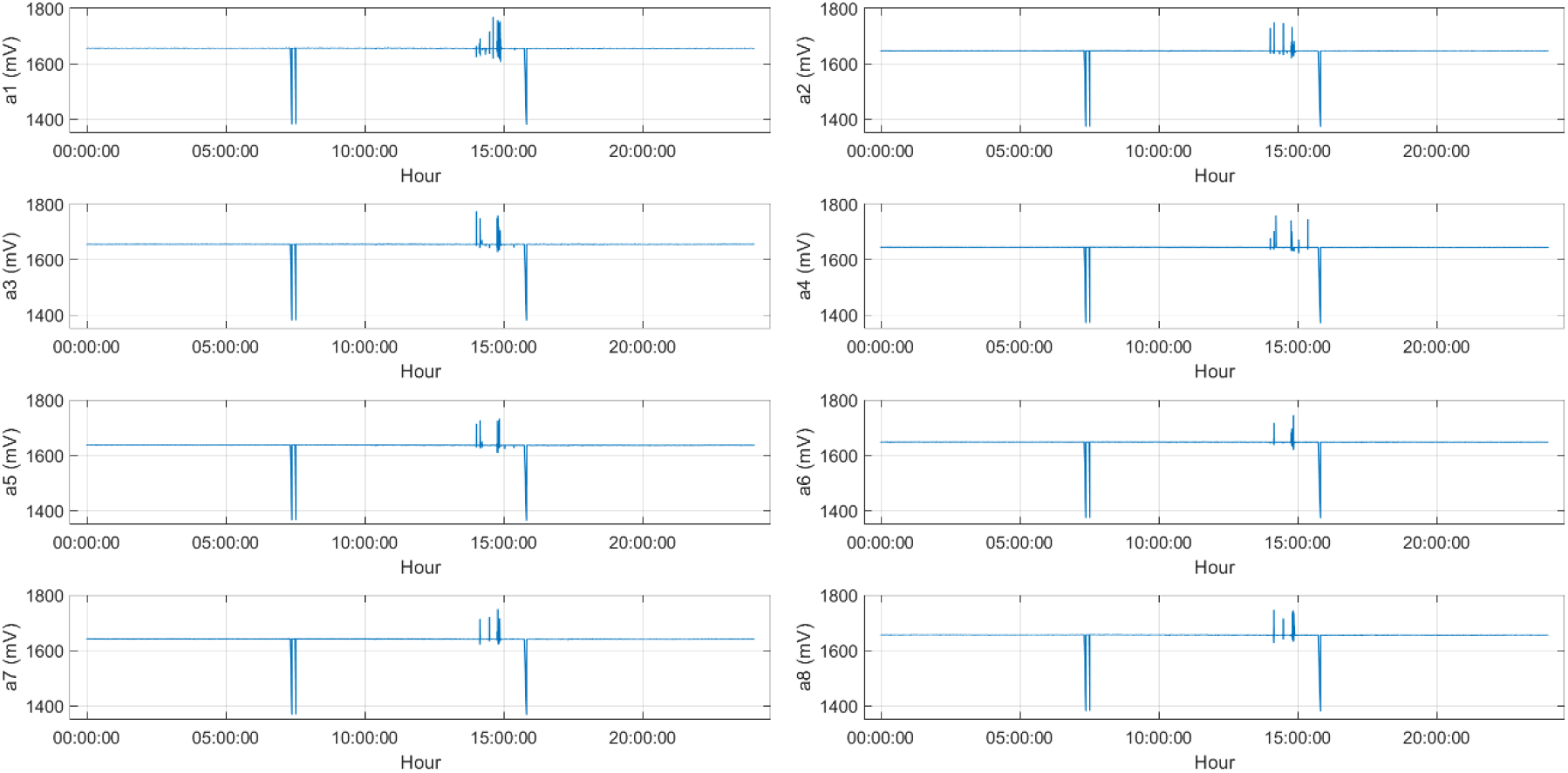
Sensor signals recorded during the day when queen bee supposedly had orientation and mating flights.

“No event” was represented by the stable sensor signal (baseline) at the level 1647 ± 7 mV, depending on sensor. The noise level of the signal was 0.45 ± 0.07 mV corresponding to 0.027 ± 0.004% of the sensor signal baseline. This indicates a very low noise level in the detection system. The data collected on the days following the queen bee mating flights were classified as “no event” signals. The event not associated with the queen detection was represented by a momentary strong drop of the signal followed by baseline restoration. The signal decrease was noted simultaneously for all sensors and it had the same magnitude (down to slightly below 1400 mV). The events of this kind were observed on several occasions during experiment. The voltage drops were caused by a sudden increase in current demand due to increased network interface card (NIC) activity, and they were eliminated by correcting the power supply circuitry. The event potentially associated with the queen detection was actually a specific sequence of up (major) and down spikes around the sensor signal baseline, occurring over some period of time. Such a sequence is visible in Fig.6 between 14:00 and 15:50. This fragment was shown in Fig. 7, zoomed.

**Fig. 7.**
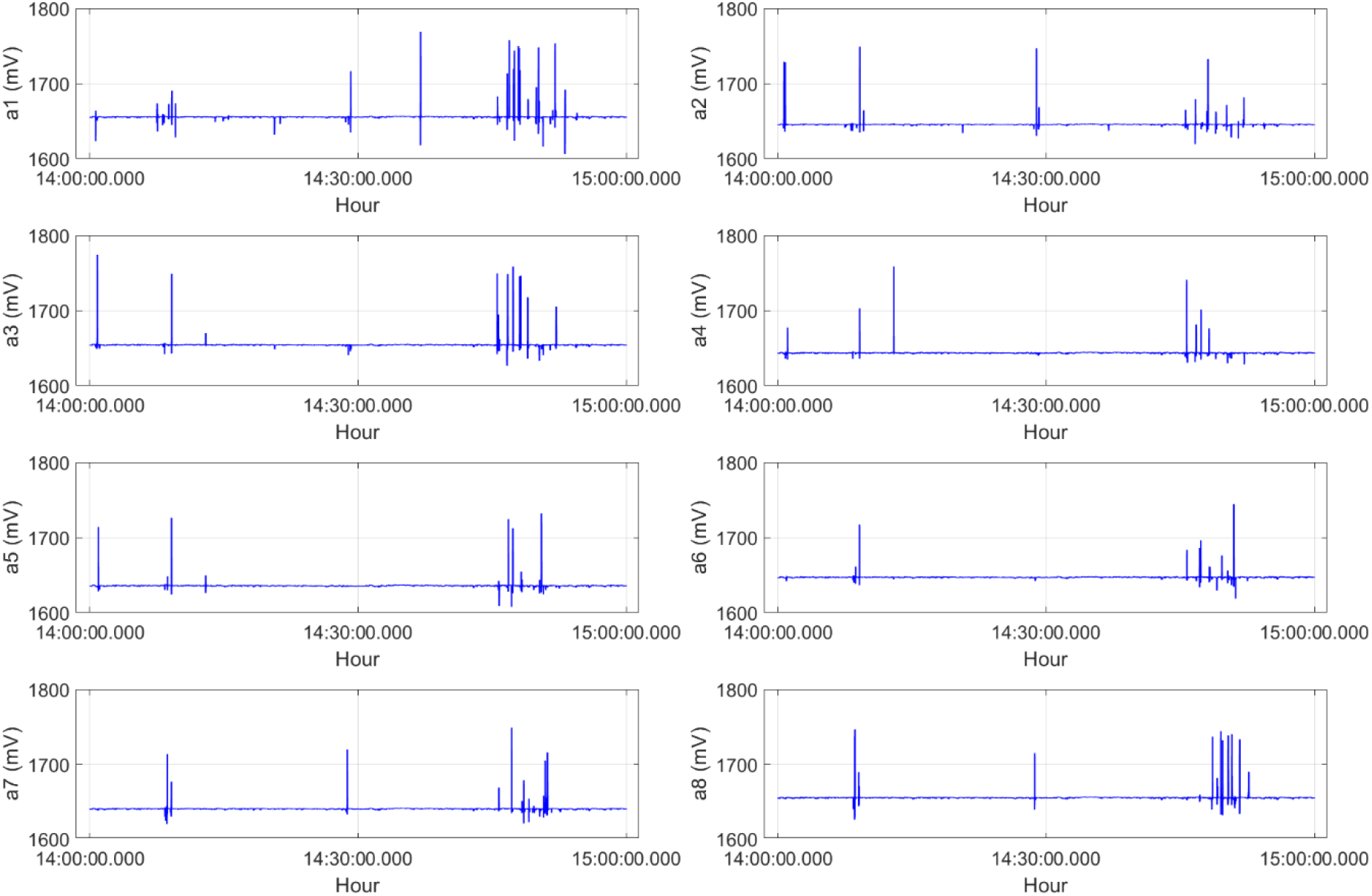
Sensor signals supposedly associated with queen bee mating flights; Zoomed time span 14:00-15:00 from Fig.6

As shown in Fig. 7, the series of spikes visible in the signals of the individual sensors were different in terms of amplitudes and time distribution. This indicates that queen bee used different regions of the hive entrance when repeatedly moving in and out. Based on Fig. 7, most spikes occurred on channels 1, 8 and 3, which correspond to the sides of the hive entrance. This observation suggests that the queen moved predominantly along the walls of the hive entrance

As shown in Fig. 7, the interpretation of the measurement data in terms of mating flight occurrence and duration is not straightforward. The results reveal quite complex and intriguing patterns of queen bee behaviour, which will be investigated in future work.

Signal to noise ratio (SNR) was determined for the detection system based on the data shown in Fig.7. The SNR was 57 ± 2dB, indicating high sensitivity of the detection system.

## 4. Strengths and limitations of the proposed system

Reliable confirmation of the presence and activity of the honey bee queen is fundamental to colony management. Under routine beekeeping conditions, this typically requires opening the hive and performing a manual inspection. Such interventions are disruptive to colony activity. They are undesirable during the queen mating period, which may extend for up to three weeks following emergence. During this time, queens perform one or more mating flights and are exposed to a substantial risk of loss before successfully returning to the colony. Consequently, a non-invasive method capable of automatically confirming queen departures and returns would represent a valuable tool for both research and practical beekeeping, especially in large-scale queen breeding.

The system presented in this work addresses this need by combining an array of Hall-effect sensors with a miniature magnetic tag attached to the queen. The feasibility of the proposed detection principle was demonstrated under both laboratory and field conditions, providing evidence that magnetic sensing constitutes a viable approach for automated queen detection and monitoring. Beyond validating the measurement concept, the experimental evaluation identified several advantages of the proposed system as well as opportunities for further refinement.

A principal strength of the system is its ability to monitor queen presence continuously and autonomously without disturbing colony activity or requiring beekeeper intervention. The achieved detection sensitivity allows reliable detection of a tagged queen moving through the hive entrance. Moreover, the spatial arrangement of multiple Hall-effect sensors provides overlapping detection regions, enabling simultaneous observation by several sensors. This redundancy increases detection reliability and reduces the probability of missed events compared with single-sensor configurations.

The developed system also offers practical advantages from an engineering perspective. It is compact, modular and portable, requiring only minimal modification of the hive structure and having negligible impact on the colony environment. The electronic architecture supports real-time notification of detection events while simultaneously recording measurement data for subsequent post-processing and behavioural analysis. Importantly, the prototype maintained stable operation throughout field experiments conducted under a range of environmental conditions, including periods of elevated temperatures, rainfall and thunderstorms, demonstrating its robustness under realistic beekeeping conditions.

Despite these encouraging results, the current implementation should be regarded as a proof-of-concept prototype, and several aspects warrant further investigation. First, the sensor array geometry may be further optimised to maximise detection coverage of the hive entrance while reducing the number of Hall-effect sensors. Such optimisation should consider both the magnetic field characteristics of the tag and the sensitivity of the Hall-effect sensors.

Second, although a commercially available miniature permanent magnet proved suitable for demonstrating the proposed concept, the development of a dedicated magnetic tag may improve overall performance. Optimisation of tag geometry, dimensions, mass, magnetic properties and attachment method could enhance detection efficiency while further reducing any potential influence on queen behaviour.

The initially applied event identification criterion based solely on a sensor signal threshold is prone to false positive detections. Based on the obtained results, queen bee mating behaviour appears to be more reliably identified from the temporal evolution of the sensor signal than from its instantaneous value. This requires careful investigation.

Another promising direction is the integration of complementary sensing modalities to provide independent confirmation of detection events. Combining magnetic sensing with additional sources of information, e.g. optical sensing could improve confidence in automated event classification.

The long-term reliability of the sensing unit may be enhanced by reducing its exposure to environmental contamination. In the current prototype, the sensor array is positioned at the hive entrance, where it remains exposed to insects, propolis deposition, beekeeper manipulation and adverse weather conditions. Future work will therefore consider optimising the placement of the sensor array within the hive entrance and developing protective solutions.

Finally, we noted several technical issues regarding battery use and limitations of the communication infrastructure, that need to be addressed.

Overall, the present study demonstrates that Hall-effect sensing represents a promising new measurement modality for non-invasive honey bee queen monitoring. Continued optimisation of both the sensing hardware and data analysis algorithms has the potential to transform the proposed system into a reliable tool for precision apiculture and for future investigations of queen behaviour and colony dynamics.

## 5. Summary and conclusions

This study presents a honey bee queen detection system based on an array of Hall-effect sensors and a magnetic tag attached to the queen.

A prototype of the system was designed, constructed, and experimentally evaluated. Its functionality was successfully demonstrated under both controlled laboratory conditions and field experiments.

Further refinement and validation of the proposed system may facilitate its application in both research and practical beekeeping.

The obtained results indicate that Hall-effect sensing provides a promising approach for continuous and non-invasive monitoring of honey bee queens.

This methodology has the potential to expand the range of available tools for colony monitoring and to support the development of advanced measurement technologies for precision apiculture.

## Acknowledgments

The research was supported by the Agency for Restructuring and Modernisation of Agriculture (ARMA) based on application No. BWI03.61835.1.1.2025, submitted under Intervention I.6.6 – Beekeeping Sector Intervention – Research and Scientific Support, within the framework of the Strategic Plan for the Common Agricultural Policy 2023–2027.

We would like to thank Mr. Stanisław Motyl for his support in conducting the field experiment, particularly for providing the experimental site, the bee colonies, and for taking care of the mating box.

